# A novel approach to comparative RNA-Seq does not support a conserved set of genes underlying animal regeneration

**DOI:** 10.1101/2021.03.22.434850

**Authors:** Noemie Sierra, Noah Olsman, Lynn Yi, Lior Pachter, Lea Goentoro, David A. Gold

## Abstract

Molecular studies of animal regeneration typically focus on conserved genes and signaling pathways that underlie morphogenesis. To date, a holistic analysis of gene expression across animals has not been attempted, as it presents a suite of problems related to differences in experimental design and gene homology. By combining orthology analysis with a novel statistical method for testing gene enrichment across large datasets, we are able to test whether biological processes across organisms share transcriptional regulation. We applied this method to six publicly available RNA-seq datasets from diverse examples of animal regeneration. We recovered 160 conserved orthologous gene clusters, which are enriched in structural genes as opposed to those regulating morphogenesis. A breakdown of gene presence/absence provides only limited support for the conservation of pathways typically implicated in regeneration, such as Wnt signaling and cell pluripotency. Specifically, these pathways are only conserved if we allow gene paralogs to be interchangeable through evolution. Overall, our analysis does not support the hypothesis that a shared set of ancestral genes underlie regeneration mechanisms in animals. The methods described in this paper will be broadly applicable for studying the genetic underpinnings of traits across distantly related organisms.

## INTRODUCTION

Why regeneration occurs in some animals and not others remains an enigma in biology. It is well known that certain groups can readily regenerate lost tissues and body parts (e.g. planarian worms, salamanders, cnidarians), while regeneration in others is restricted to specific organs or developmental stages (e.g. nematode worms, insects, mammals). Animals with strong regenerative capabilities are distributed across the evolutionary tree without a clear pattern (1), and even closely related species can demonstrate dramatically different capacities (2, 3). These observations lead to two competing evolutionary scenarios: body regeneration is either an ancient, conserved animal trait that has been lost to varying degrees across multiple lineages, or it is a derived trait that multiple lineages have converged upon independently. Resolving these competing hypotheses has profound consequences for the goals of comparative regenerative biology: are we searching for unifying principles, or trying to determine how various animals deal with the universal problem of body damage?

While many studies focus on putative candidate genes underlying animal regeneration, a growing body of literature challenges any simplistic interpretation. Some genes and pathways occur commonly in studies: Wnt signaling, for example, offers a compelling candidate for a “master regulator” of stem cell dynamics during regeneration (4), as it has been shown to play a critical role in planarian worms (5, 6), fish (7), amphibians (8), and mammals(9–11). In contrast, several recent studies suggest that key components of regeneration might be dissimilar across major groups. For example, a MARCKS-like protein that initiates limb regeneration in axolotl salamanders appears to be a vertebrate novelty (12). Regeneration in newts, a different group of amphibians, involves genes not found in the axolotl (13). Finally, genes such as the Oct4/POU5F1 “master regulator” of stem cell pluripotency appear absent in invertebrates (14). It is unclear whether these observations represent anomalies obfuscating a conserved set of shared genes, or if they hint at the true evolutionary convergence driving animal regeneration.

Whether the molecular mechanisms of regeneration are conserved across animals rests, in part, on what counts as a “conserved” (i.e. homologous) gene. Homologous genes can be subdivided into orthologs—genes related by vertical descent from a common ancestor—and paralogs—genes that arise by duplication events. Orthologs or paralogs may perform similar functions, but in evolutionary biology, common ancestry is what defines conservation. Paralogs by definition cannot be traced back to a single gene in a last common ancestor, and the utilization of paralogs by different species, even in similar biological processes, generally implies evolutionary convergence. Complicating this matter, the ortholog/paralog distinction is contingent on the organisms being studied. As more distantly related species are analyzed, families of paralogous genes often collapse into a single orthologous clade (see **Figure S1** for an example). Tests of molecular conservation therefore require careful consideration of the evolutionary history of genes.

**Figure S1.**
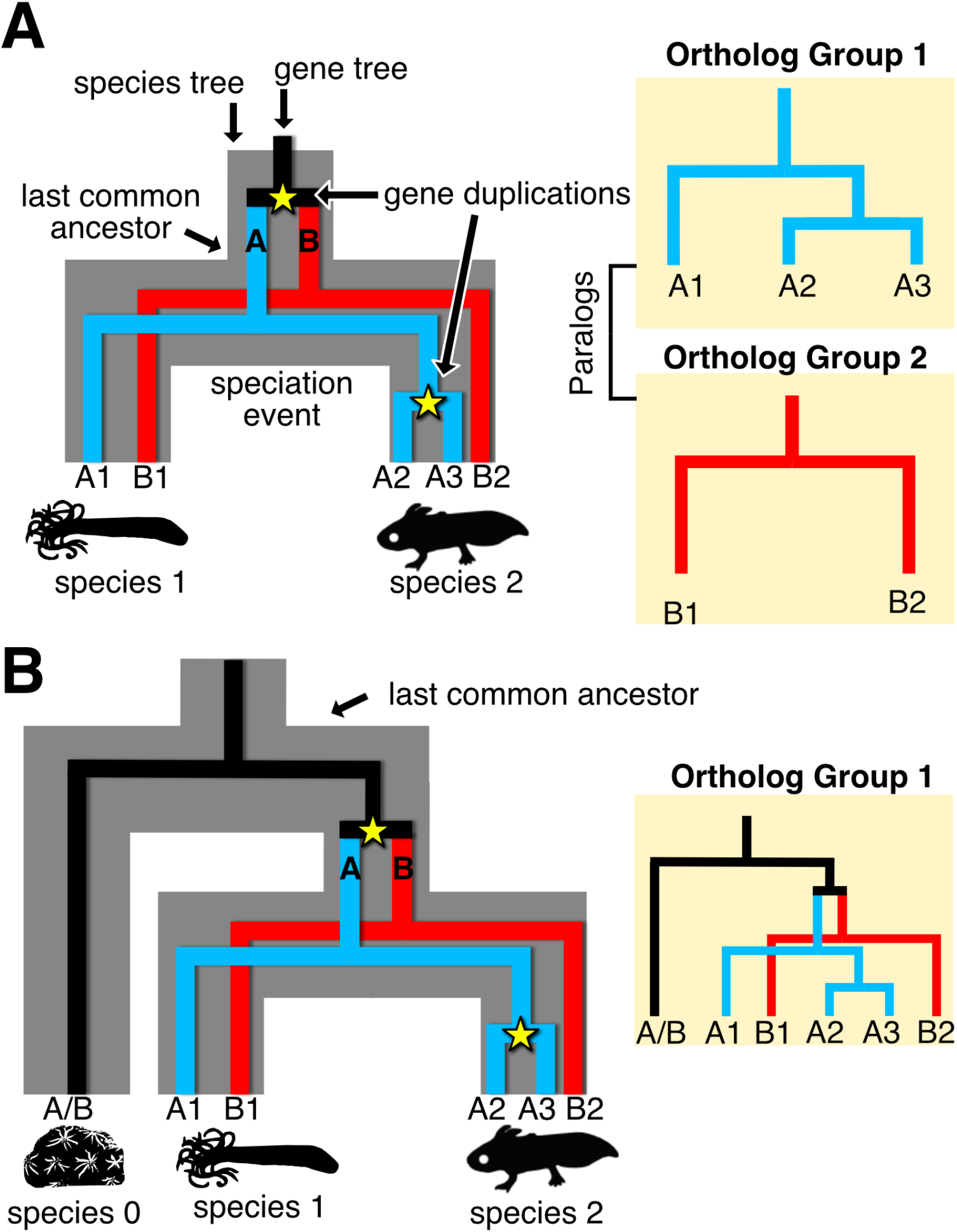
Evolutionary history and the ortholog / paralog distinction. (**A**) A hypothetical gene family and its evolution in two species. In this scenario, a duplication event occurred before the split of species 1 and species 2, leading to paralogs “A” and “B”. As two genes were present in the last common ancestor, the genes can be separated into two discreet conserved orthologous groups (COGs). (**B**) The same scenario as (A) with an additional species included. In this scenario all genes in the three living species can be traced back to a single gene in the last common ancestor. From this evolutionary vantage, all 6 genes collapse into one COG.

The problem described above is compounded when using RNA-seq technology to identify “conserved” genes between distantly related taxa undergoing similar biological processes. There are two major hurdles in comparing datasets across distantly related animals. Firstly, genes rarely share one-to-one homology across species. An ancestral gene might, over the course of evolution, undergo multiple rounds of duplication, resulting in a single gene in species 1, two homologs in species 2, and eight homologs in species 3. Secondly, RNA-seq studies have varying temporal resolutions, timescales, and depths of sequencing. These two issues result in a heterogeneous list of statistical tests that are problematic to compare between studies. As an example, imagine a conserved orthologous gene group, where species 1 has one gene sampled at three time points, while species 2 has five paralogous genes sampled at seven time points. If all time points are compared to each other, this would result in six statistical tests for species 1 compared to 25,200 tests for species 2.

To address this discrepancy, we used a Lancaster p-value aggregation method, which provides a systematic way of collapsing multiple statistical tests from RNA-seq studies into one value (15, 16). In our case, the multiple tests include time sampling of all genes that are members of a conserved ortholog group (COG). The method looks at the p-values generated from adjacent time points in a differential gene expression analysis, and treats each as an independent significance test of the hypothesis that the broader COG is differentially expressed. This aggregation method from (16) takes advantage of the fact that many independent p-values generated by the null hypothesis should follow a uniform distribution on the interval (0,1). Consequently, we can test the *uniformity* of the set of p-values to determine their likelihood of being generated from the null hypothesis, which in our case corresponds to the COG not being differentially expressed during regeneration. This method will capture all genes that would pass a significance test in a standard RNA-seq analysis, as well as COGs that have more borderline-significant p-values than would be expected by chance. This approach allows us to identify relevant gene families that could be missed in a typical RNA-seq analysis, as well as make statistically honest comparisons of differential gene expression between diverse studies.

In this study, we compared publicly available RNA-seq datasets spanning wildly different organisms and structures undergoing regeneration (**Figure 1A**) to determine if an underlying core set of genes could be elucidated. The datasets analyzed include tissue regeneration in sea sponges (17), oral/aboral body regeneration in sea anemones (18), head/tail regeneration in planarian worms (19), regeneration of “Cuvierian tubules” in the respiratory system of sea cucumbers (20), hair cell regeneration in zebrafish (21), and limb regeneration in axolotl salamanders (22). These datasets are highly divergent in their sampling regimes but cover the relevant early window between wound healing and blastema formation/ cell proliferation (**Figure 1A**). Our approach involved clustering all proteins from all six species into sets of COGs. We then performed p-value aggregation on each RNA-seq dataset (i.e. species), reducing the many p-values from multiple orthologs and time points into a single 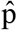-value for each COG. The 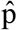-values for each COG were compared between datasets to determine which COGs were differentially expressed across all six species. This procedure is graphically illustrated in **Figure 1B**, with a more in-depth flow chart provided in **Figure S2**. Despite the limitations inherent in comparative RNA-seq (considered in detail in the Discussion), this study provides a first-order analysis to clarify what is conserved in animal regeneration at a molecular level.

**Figure 1.**
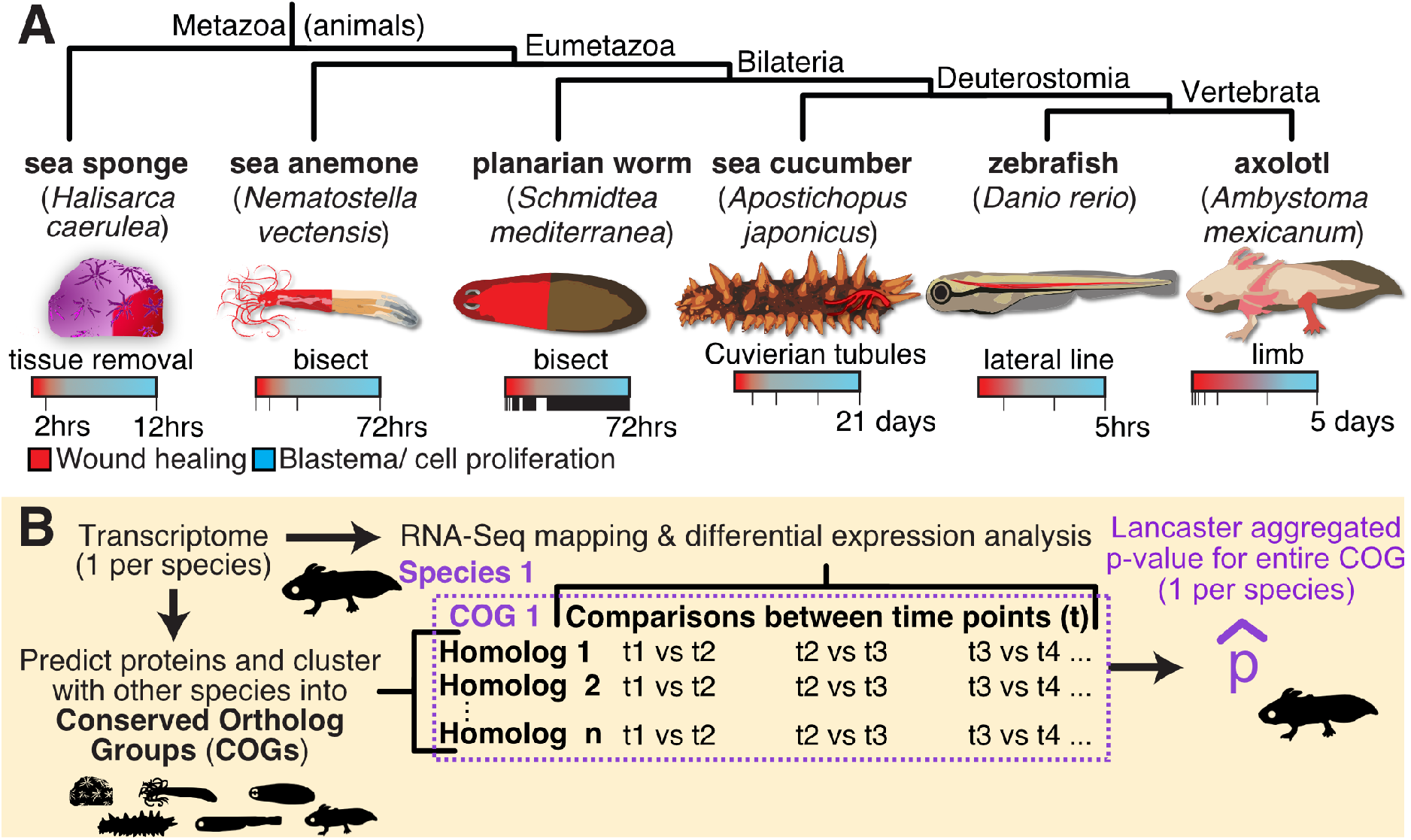
Cases of animal regeneration included in this study. (**A**) The six animals analyzed in this paper, organized by their evolutionary relationships. The region of each organism undergoing regeneration is highlighted in red and is described underneath the image of each animal. The RNA-seq sampling regime from each study is visualized with a bar; each time point that was sampled is represented by a notch in that bar. Despite the different absolute time ranges, the studies are comparable in that they all analyze early key stages of regeneration: starting with wound healing (red) and transitioning into blastema formation / cell proliferation (blue). (**B**) Graphical summary of the approach used to compare RNA-seq data between the six datasets. A more detailed protocol is visualized in Figure S2.

**Figure S2.**
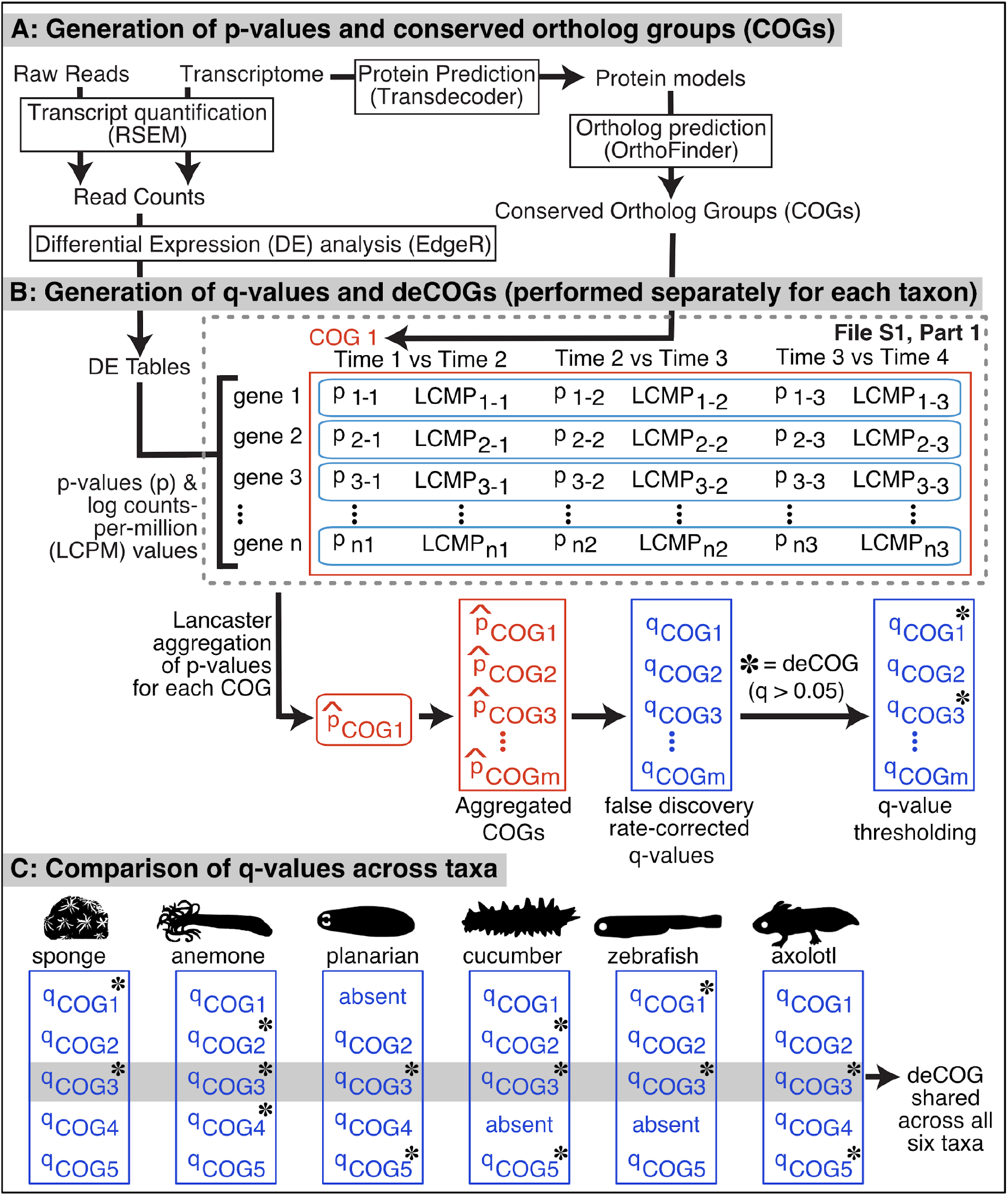
Graphical Overview of methodology for identifying differentially expressed conserved ortholog groups (deCOGs).

## RESULTS

The first step was to organize all genes from our six species into clusters of putative orthologs. We used OrthoFinder (23) to assign orthology, as this program combines amino acid sequence similarity and phylogenetic relationships to reconstruct the evolutionary history of gene families. OrthoFinder assigned 266,324 proteins generated from the six reference transcriptomes into 16,116 conserved orthologous groups or “COGs” (see **Table S1** for detailed OrthoFinder results). These COGs were typically large, with a mean of 16.5 genes per COG. This reflects the large number of gene models in certain datasets (particularly the axolotl and zebrafish) as well as the wide evolutionary vantage taken in this study. Because we assigned orthology at the pan-animal scale, many paralogs in vertebrates or eumetazoans collapsed into a single COG in this study (see **Figure S1** for an example). After genes were assigned to COGs, we used the Lancaster method to aggregate all p-values per dataset per COG into one 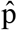-value (16). If that 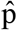-value met a false-discovery adjusted threshold of 0.05, we considered the COG differentially expressed for that particular dataset.

We recovered 160 COGs that were differentially expressed across all six species’ RNA-Seq datasets, which we treat as a generous estimation of genetic conservation. There are multiple reasons this number likely overestimates the amount of conservation between datasets. Firstly, we are not considering when a gene is being expressed (e.g. wound healing versus blastema formation) nor are we considering the direction of gene expression (e.g. upregulation versus downregulation). We are also grouping genes into animal-wide COGs that, with further inspection, can be typically divided into distinct paralogs at finer evolutionary scales (a point we consider later in the text). Alternatively, there are several counterarguments suggesting 160 COGs could be an undercount. The quality of one or more datasets could result in us missing differentially expressed COGs. For example, the sea cucumber *A. japonicus* has a limited RNA-Seq dataset and no reference genome; it is possible that some COGs are missing from our list because the relevant homologous gene(s) were not reconstructed from the *A. japonicus* dataset, or there was not enough data to get statistical support for differential expression from this species. It is therefore important to test whether removing one or more datasets dramatically increases the number of COGs we recover.

To test how robust the assignment of differentially expressed COGs (deCOGs) is to differences between datasets, we examined how adding and removing datasets impacted the final number of deCOGs (illustrated in **Figure S3**). Removing any particular dataset from the study increased the number of deCOGs shared across the remaining 5 datasets by an additional 26 to 196, depending on which dataset was removed. In other words, considering any 5 of the 6 datasets recovers a comparable number of deCOGS. Moreover, we did not find any correlation between the quality of the RNA-Seq study and the number of additional deCOGs recovered when a dataset was removed. For example, removing the sea anemone from the analysis provided the greatest increase in deCOGs, even though this high-quality dataset included four RNA-Seq time points with biological replicates, as well as a well-annotated genome to work off of. Conversely, the sea sponge had the poorest sampling regime, yet its removal resulted in one of the smallest gains (+49 deCOGs). While some deCOGs could be lost due to incomplete sampling of gene expression during regeneration, our analyses do not suggest an obvious bias caused by the quality of the datasets under consideration.

**Figure S3.**
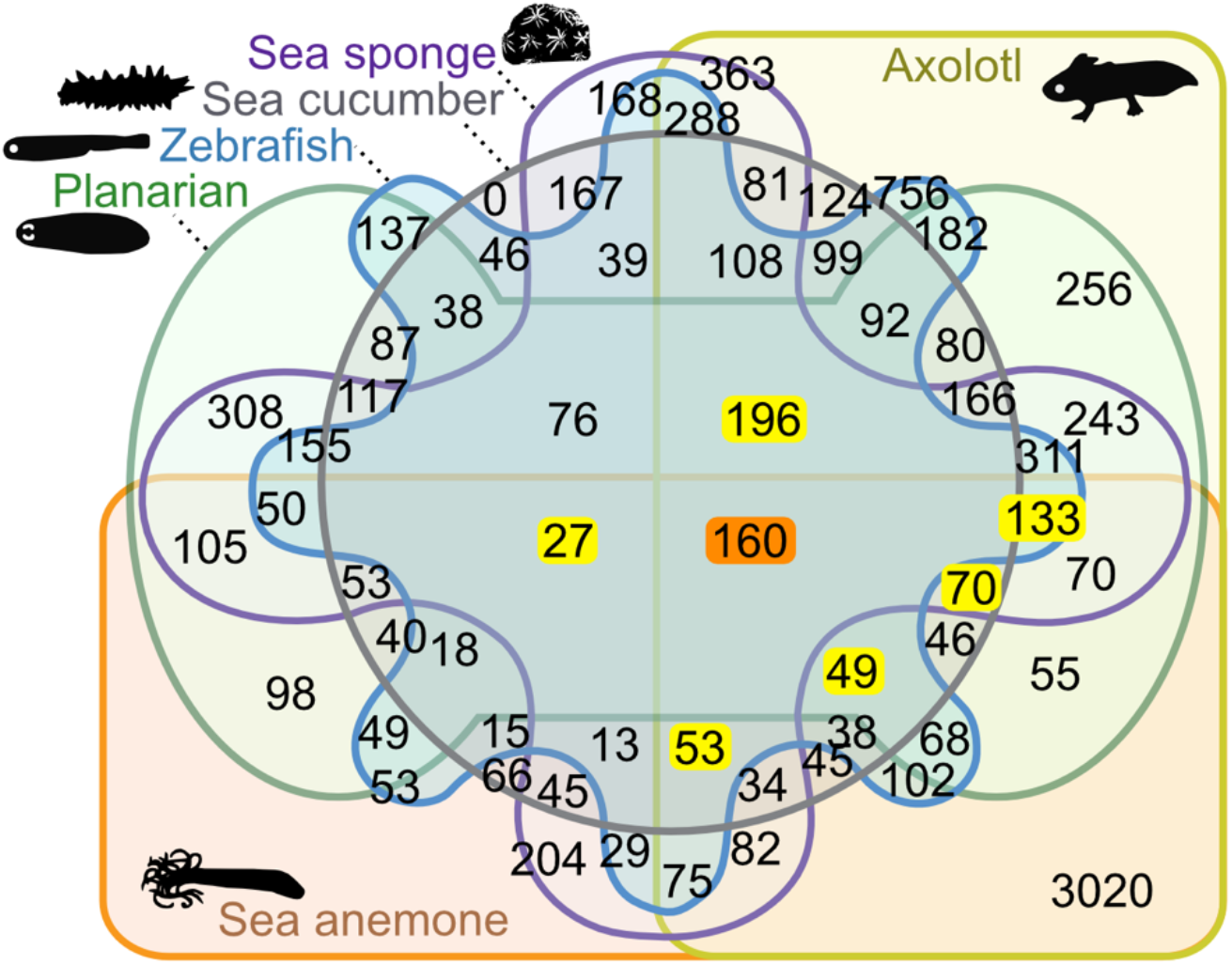
An Edwards-Venn diagram demonstrating the number of overlapping differentially expressed conserved orthologous groups (deCOGs) across all 6 datasets. The number of deCOGs common across all 6 cases (160) is highlighted in orange. Additional deCOGs that are recovered when individual case studies are removed are highlighted in yellow.

Following this check on the data, we proceeded with a holistic assay of COGs to see how similar the datasets were to each other. We used presence/absence data to construct a correlation matrix that illustrates the total number of COGs shared across datasets (**Figure 2A**) and a second matrix restricted to COGs that are differentially expressed in one or more datasets (**Figure 2B**). Our results suggest the datasets exhibit dramatically distinct gene repertoires. Both matrices organize the taxa on evolutionary relationships—albeit imperfectly—with vertebrates forming one major clade and the paraphyletic invertebrates forming a second. The similarity between vertebrate datasets (axolotl and zebrafish) is surprising, since the structures being regenerated are so divergent—limb versus a cell type respectively. This suggests evolutionary relationship is more predictive of transcriptional similarity than the type of structure that’s regenerating. Additionally, if genes expressed during regeneration represented an evolutionarily conserved network, we would anticipate the deCOG correlation matrix in **Figure 2B** to show greater similarity than the full COG matrix in **Figure 2A**. Instead, the correlation matrix looks fairly similar, and even shows less similarities between the invertebrates. This suggests that genes expressed during regeneration are no more similar across datasets than expected by chance.

**Figure 2.**
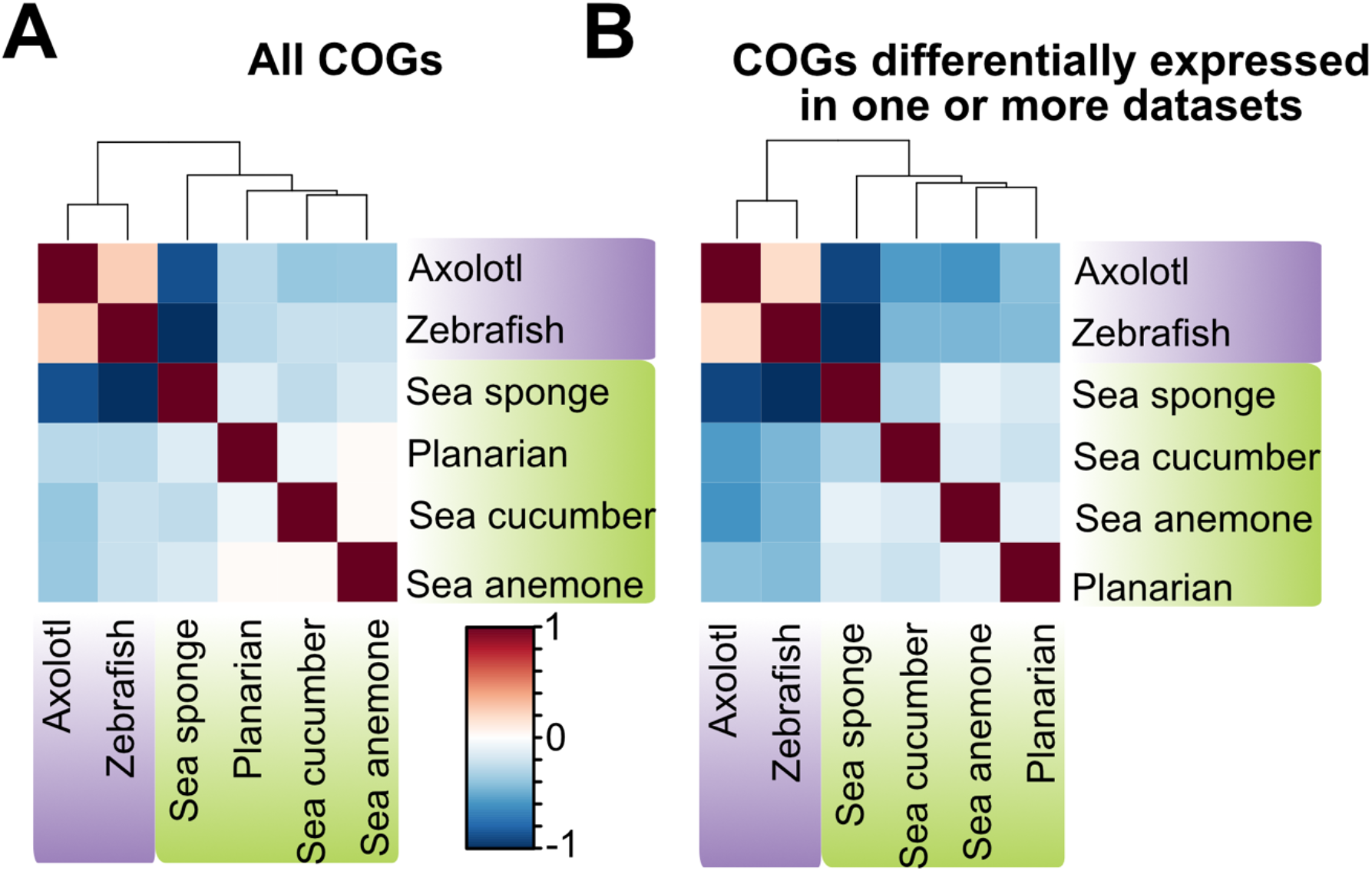
Correlation matrices based on the presence/absence of COGs across taxa. The scale is calculated using the Pearson correlation coefficient, where −1 indicates a perfectly negative linear correlation between two variables and 1 indicates a perfectly positive linear correlation between two variables. (A) Matrix derived from all COGs as assigned by OrthoFinder. (B) The same analysis restricted to differentially expressed COGs (deCOGs).

One of the patterns seen in **Figure 2** (and see **Figure S3**) is that the vertebrates (the axolotl and zebrafish) appear more similar to each other than any other combination of taxa. This raises the possibility that regeneration in vertebrates is driven by shared, vertebrate-specific genes. To test this hypothesis, we calculated how many deCOGs exist at each node of the evolutionary tree (**Figure 3**), starting with the 160 deCOGs shared across all animals and then seeing how many additional COGs are recovered when we restrict our analyses to more derived evolutionary clades. We then assigned all of these deCOGs a “phyletic origin” by comparing the protein models to those in NCBI (see Methods for details). The results, shown in **Figure 3**, suggest that the majority of deCOGs have pre-metazoan, eukaryotic origins. In other words, regeneration in most animal groups does not appear to require much input from novel, metazoan-specific genes. While this trend holds true in vertebrates, ~9% of all deCOGs unique to this clade do appear to be vertebrate-specific novelties. This suggests that while the genetic control of animal regeneration is largely driven by the co-option of ancient, pre-metazoan genes, regeneration in vertebrates also requires input from genes unique to the group.

**Figure 3.**
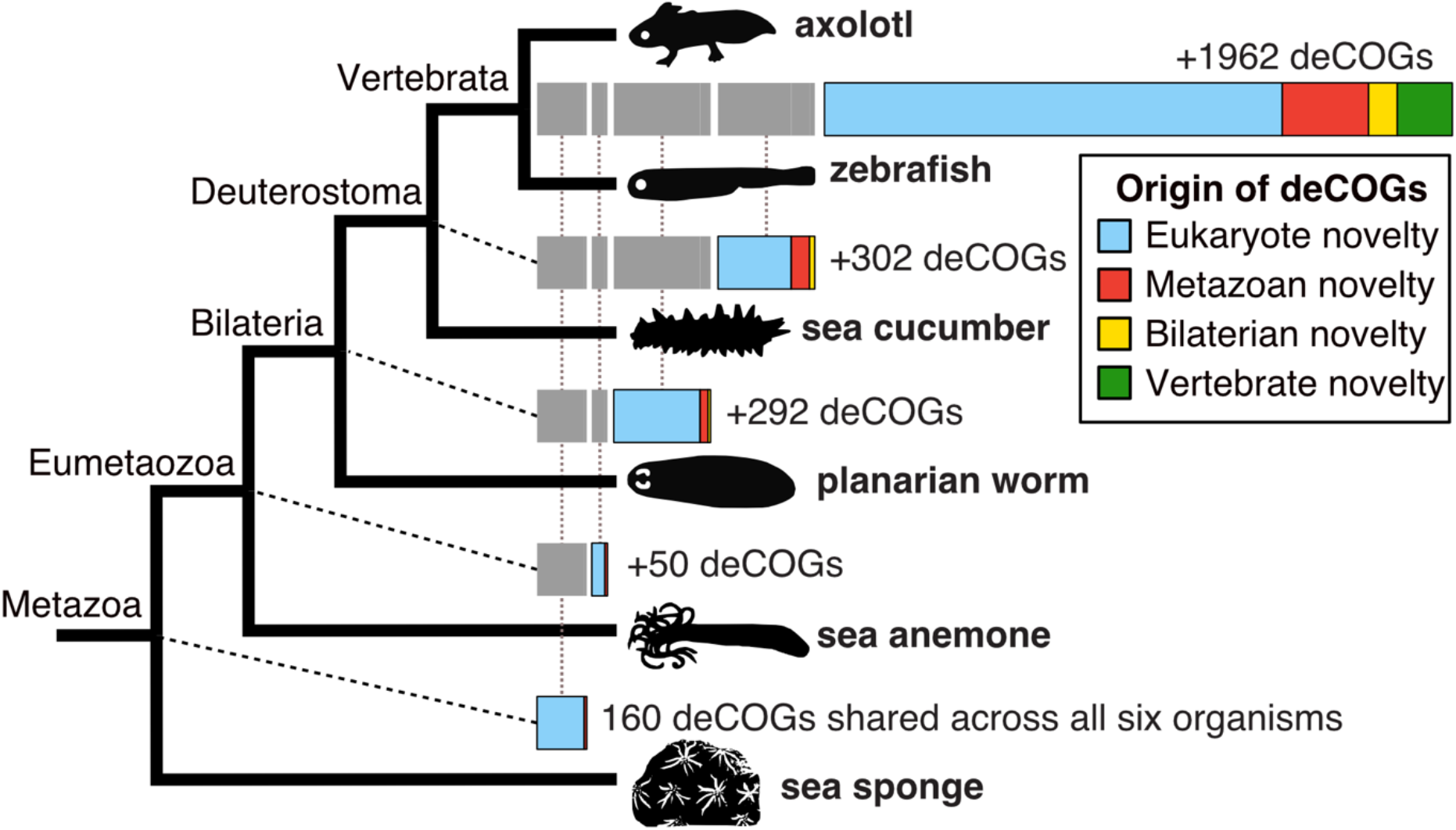
Evolutionary (phyletic) origin of deCOGs. The total number of deCOGs recovered at each node of the evolutionary tree is indicated by a bar chart to the right. Novel deCOGs at each node are broken down by their phyletic origin; for example, deCOGs that are a “bilaterian novelty” contain genes that have no significant sequence similarity to genes outside of the Bilateria.

To explore the possible function of the 160 deCOGs recovered across all taxa, we used two highly-cited web resources, STRING (24) and DAVID (25), to perform functional enrichment analysis. We focused on the zebrafish for these analyses, as it represents the best-studied organism in our dataset. The 160 deCOGs include 2,182 zebrafish transcripts, 554 of which could be considered differentially expressed (using the generous cutoff of raw p-values < 0.01). We compared this list of genes against the zebrafish genome to look for enriched biological pathways using the comprehensive and highly-cited Kyoto Encyclopedia of Genes and Genomes (KEGG) database (see Additional File 1, part 4 for full results). According to STRING and DAVID analyses, the 554 differentially expressed zebrafish genes are enriched in basic cell processes, including melanogenesis, regulation of the actin cytoskeleton, phagosomes, and focal adhesion (see **Tables 1 and 2**). Regarding KEGG pathways, Notch signaling is recovered in both analyses, while Wnt, FoxO, and mTOR pathways are enriched in the STRING analysis. Unfortunately, deeper study of the genes driving “enrichment” challenge the hypothesis that these pathways are being utilized across all species. In all of these pathways, enrichment is primarily driven by multiple paralogs of the same few genes being differentially expressed in the zebrafish. For example, Wnt and Frizzled paralogs represent 9 out of 11 genes driving Wnt enrichment and 9 of the 15 genes driving mTOR enrichment. It is worth noting that Wnt and Frizzled are not generally cited as part of the canonical mTOR pathway, and none of the proteins in mTOR Complex 1 or 2 were recovered from this dataset (see **File S1, part 4.1** for the list of genes). Similarly, Notch enrichment is driven by the presence of 8 differentially expressed genes, 7 of which are Delta/Jagged paralogs. If these pathways were truly playing a conserved role in regeneration, we would anticipate more genes in these pathways being differentially expressed across all datasets. Re-running the analysis with an expanded list of deCOGs based on evolutionary subclades (see **Figure 3**) did not have a major impact on the pathways recovered. However, when we restricted our analysis to deCOGs shared between the vertebrates, we found a dramatic increase in the number of Wnt pathway genes represented (58 genes). Furthermore, FoxO (65 genes) and p53 signaling (32 genes) were also recovered as significantly overrepresented pathways. All of these pathways have been implicated in vertebrate regeneration (26–29). These results further support the hypothesis that a conserved regeneration network might exist across vertebrates, even though there is little evidence for conservation across the animals as a whole.

**Table 1:** Functional Enrichment of the differentially expressed zebrafish genes from our 160 deCOGs, based on DAVID.

| Term | count | % | p-value | benjamini |
| --- | --- | --- | --- | --- |
| Melanogenesis | 15 | 3.3 | 9.40E-05 | 8.40E-03 |
| Oocyte meiosis | 14 | 3.1 | 3.50E-04 | 1.60E-02 |
| Adrenergic signaling in cardiomyocytes | 16 | 3.6 | 7.70E-04 | 2.30E-02 |
| Protein processing in endoplasmic reticulum | 15 | 3.3 | 1.60E-03 | 3.60E-02 |
| Regulation of actin cytoskeleton | 19 | 4.2 | 2.10E-03 | 3.70E-02 |
| Notch signaling pathway | 8 | 1.8 | 2.20E-03 | 3.20E-02 |
| Phagosome | 13 | 2.9 | 3.30E-03 | 4.20E-02 |
| Calcium signaling pathway | 18 | 4 | 3.30E-03 | 3.70E-02 |
| Focal adhesion | 17 | 3.8 | 4.90E-03 | 4.80E-02 |

**Table 2:** Functional Enrichment of the differentially expressed zebrafish genes from our 160 deCOGs, based on STRING.

| Term | observed gene count | background gene count | false discovery rate |
| --- | --- | --- | --- |
| Phagosome | 15 | 142 | 1.41E-06 |
| Regulation of actin cytoskeleton | 19 | 251 | 1.41E-06 |
| Apoptosis | 15 | 159 | 1.88E-06 |
| Focal adhesion | 18 | 236 | 1.88E-06 |
| Protein processing in endoplasmic reticulum | 15 | 176 | 3.83E-06 |
| mTOR signaling pathway | 15 | 179 | 3.91E-06 |
| Tight junction | 14 | 203 | 7.15E-05 |
| Notch signaling pathway | 8 | 60 | 9.07E-05 |
| Oocyte meiosis | 11 | 130 | 9.60E-05 |
| Adherens junction | 9 | 95 | 0.00025 |
| Melanogenesis | 10 | 128 | 0.00038 |
| Wnt signaling pathway | 11 | 170 | 0.00072 |
| Necroptosis | 10 | 152 | 0.0011 |
| Adrenergic signaling in cardiomyocytes | 11 | 180 | 0.0011 |
| Gap junction | 9 | 123 | 0.0011 |
| Cell cycle | 9 | 137 | 0.0021 |
| Cardiac muscle contraction | 7 | 89 | 0.0033 |
| Lysine degradation | 6 | 72 | 0.0058 |
| Calcium signaling pathway | 11 | 245 | 0.0085 |
| ECM-receptor interaction | 6 | 82 | 0.0096 |
| Endocytosis | 12 | 293 | 0.0101 |
| AGE-RAGE signaling pathway in diabetic complications | 7 | 119 | 0.0123 |
| Autophagy - animal | 8 | 155 | 0.0131 |
| ABC transporters | 4 | 39 | 0.0149 |
| FoxO signaling pathway | 8 | 161 | 0.0149 |
| Ferroptosis | 4 | 42 | 0.0175 |
| Cell adhesion molecules (CAMs) | 6 | 120 | 0.0407 |

The enrichment analyses described above demonstrate the importance of distinguishing clade-specific patterns of orthologs and paralogs; when we dig into the data, it becomes clear that many of our deCOGs are driven by the differential expression of evolutionary paralogs. A good example of this issue comes from the Wnt family of genes, which are recovered as a single deCOG in our analysis. The gene tree produced by OrthoFinder is reprinted in **Figure 4**. Our analysis suggests that sponge Wnt genes cannot be assigned to the known subfamilies of “higher” animals, resulting in all Wnts collapsing into one COG (see Borisenko et. al (30) for similar results). Ignoring the sponge, only one of the Wnt subfamilies (Wnt8/9) is present in all organisms in our analysis, and no Wnt subfamily demonstrates differential expression across all taxa. So, while Wnt genes are differentially expressed in every example of regeneration, each organism uses a different combination of paralogs. This result could be interpreted as evidence that diverse Wnt genes can be removed and integrated into a conserved regeneration gene network, or alternatively, that different organisms have independently integrated Wnt signaling into regeneration. Either way, this case study illustrates that a deCOG is not synonymous with a conserved gene, and offers no support that an ancestral Wnt protein has a conserved function in regeneration across animal evolution.

**Figure 4.**
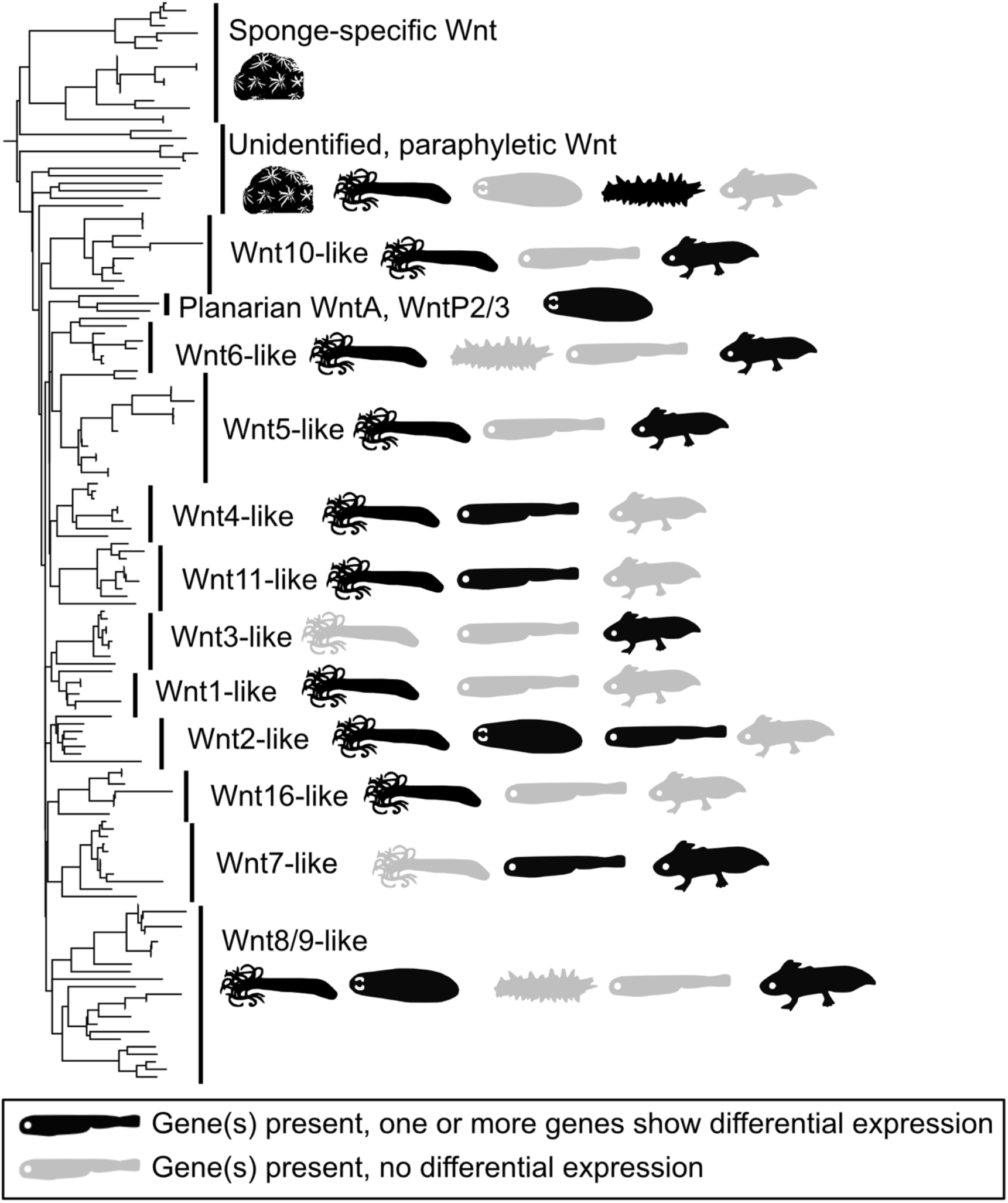
The presence of Wnt genes in the 6 RNA-Seq datasets analyzed (produced by OrthoFinder). Wnt genes were recovered as a single deCOG in our analysis, which we manually subdivided into 13 previously described subfamilies. The presence/absence of these subfamilies in each taxon is demonstrated by silhouettes. Grey silhouettes show the subfamily is present in the organism’s transcriptome; black silhouettes show that the subfamily is present and differentially expressed in the relevant RNA-Seq study. Note that no subfamily is present and differentially expressed across all taxa.

Given the longstanding interest in stem cell dynamics as a critical regulator in animal regeneration, we decided to conclude our analysis by exploring the representation of relevant pathway in our data. **Figure 5** presents a simplified version of the KEGG stem cell pluripotency network (KEGG 04550), colored to indicate the number of datasets with one or more differentially expressed genes from the relevant COG. Few molecular signaling components were differentially expressed across all 6 datasets, and most downstream signaling targets were expressed in fewer than four datasets. Additionally, the ultimate target of these pathways—the core transcriptional network driving mammalian stem cell pluripotency (31)—were largely absent, with two of the genes missing from all datasets (Oct4/POU5F1 and Nanog). At first glance, some interesting signaling and receptor proteins appeared to be conserved across all six taxa. However, detailed analysis of the relevant COGs revealed that every example involves clade-specific paralogs being collapsed into a single pan-metazoan COG, as described previously for Wnt. Examples include “Activin” and “BMP4” being part of a single deCOG that also contains BMP2/4/5/6/8/15/16, and the “SOX2” deCOG that also contains SOX1/3/9/14 (see **Table S2, Figure S4**, and **Additional File 1, part 7** for details). We therefore find limited support for conserved genes in the cell pluripotency network, and find “conservation” in a few pathways only in the context of equating paralogs across various evolutionary lineages.

**Figure 5.**
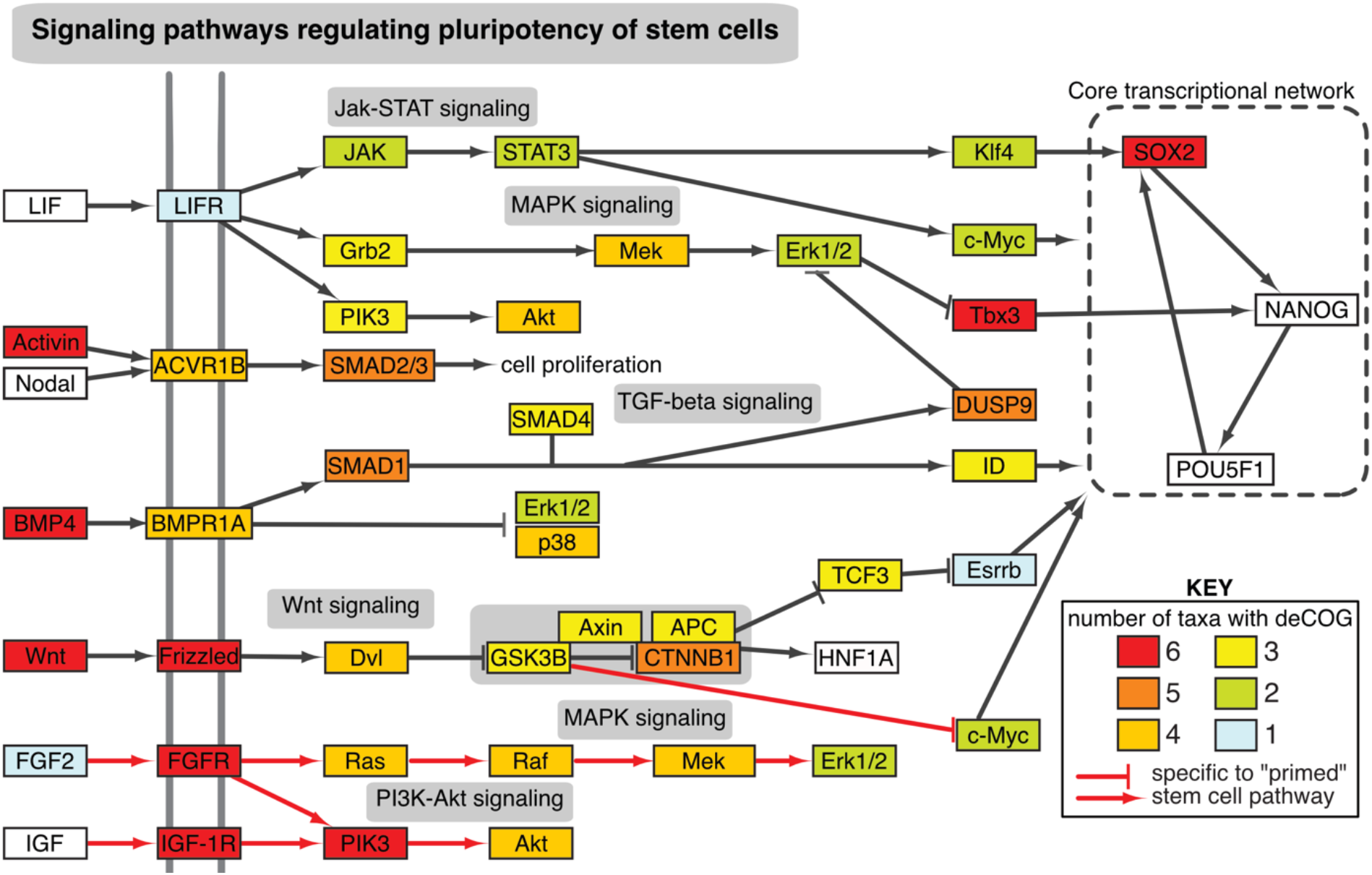
The presence of deCOGs within the stem cell pluripotency network. The network has been reproduced and simplified from KEGG pathway 04550. The color of each box indicates the number of datasets with one or more differentially expressed genes within the relevant COG. Red arrows indicate pathways that are specific to “primed” stem cells (e.g. human embryonic stem cells, human induced pluripotent stem cells, mouse epiblast-derived stem cells), grey arrows indicate pathways also found in “naïve” stem cells (e.g. mouse embryonic stem cells, mouse induced pluripotent stem cells).

**Figure S4.**
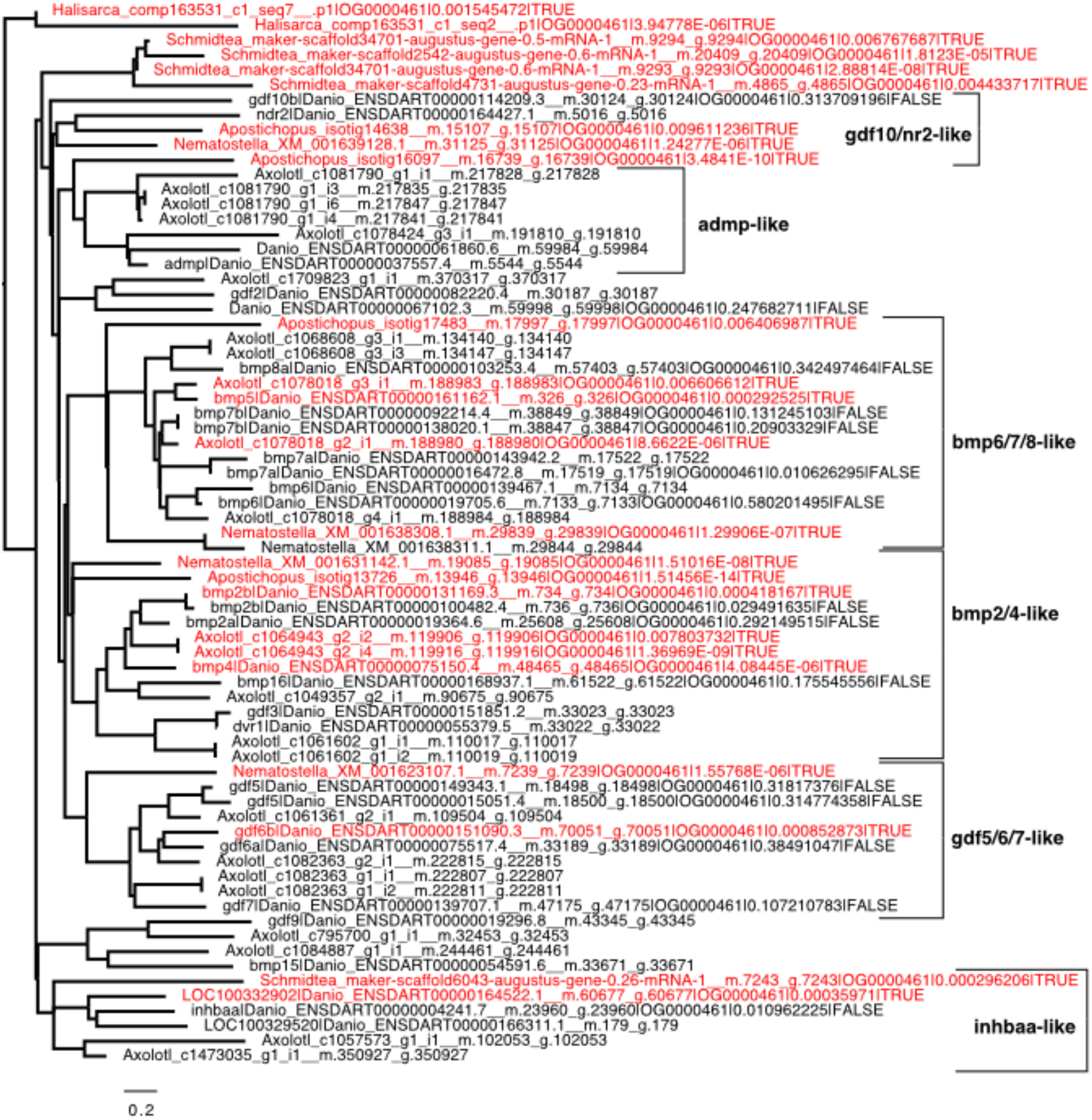
Gene tree for COG “OG0000461,” which includes activin and bmp4 genes. Genes considered differentially expressed are labeled red. Trees for the other COGs from **Figure 5** are provided in Additional File 1, part 7.

## DISCUSSION

In this study, we have found little evidence for a shared “core” network of orthologous genes across six RNA-Seq studies of animal regeneration. There are several ways to interpret our results. One possibility is that a shared genetic network underlies animal regeneration, but we failed to recover it because of the limitations of RNA-Seq. There are several arguments suggesting that this is unlikely. First, while it is true that the six datasets included in this study had markedly different sampling regimes (**Figure 1**), all of them capture the important early stages of regeneration (wound healing, blastema formation, and cell proliferation). Second, removing any single taxon had minimal impact on our results. Finally, the fact that phylogenetic relatedness is more predictive of gene content than the RNA sampling regime (**Figure 2**) or the type of regeneration that is occurring, suggests that sampling variation cannot explain the differences in gene expression. So although we cannot reject the hypothesis that deeper sampling could increase the number of shared genes, we feel confident that our results reflect a lack of a shared group of core “regeneration genes”.

A second interpretation of our results is that the shared genes we did discover play currently under-appreciated roles in driving regeneration. The 160 deCOGs recovered across all six taxa are enriched in cellular processes such as adhesion and actin cytoskeleton regulation, rather than regulatory signaling pathways or transcription factors driving morphogenesis (**Table 1**). Cellular and tissue dynamics could be critically important to initiating and maintaining regeneration. For instance, the role of musculature in driving regeneration—perhaps by providing axial specification to blastema stem cells—is supported by several studies. For example, actin-driven mechanical forces are required for the regeneration of mammalian skin (32). A focus on these genes in regeneration models might reveal conserved mechanisms driven by cell dynamics.

A third possibility is that regeneration is not a conserved process across animals at the transcriptional level. The regulatory mechanisms driving regeneration in vertebrates may not be the same as those in planarians, which in turn may be unique from those in sea anemones and other early-branching animal lineages. The conservation of cellular processes in our “core” gene list could simply be a byproduct of basic cellular necessities; for instance, actin movement being necessary for wound closure. Similarly, the presence of many Wnt signaling genes across our datasets (and across studies of regeneration more broadly) could simply reflect the fact that there are a limited number of cell signaling pathways that animals use to pattern tissues. The lack of conservation in Wnt paralog usage or downstream pathway targets supports this hypothesis. It is worth reiterating that our analysis was designed to err on the side of being overly inclusive; we treated all genes as “differentially expressed” regardless of *when* they were expressed, or whether the genes were up- or down-regulated. This further challenges the limited examples of conservation we recovered. As an example, one of the major conclusions from the original zebrafish RNA-seq study was that Wnt signaling is upregulated hours after the onset of stem cell proliferation, which is in contrast to expectations based off of other model systems where it is typically downregulated (21). Given our forgiving analysis design, combined with the fact that each dataset includes hundreds to thousands of differentially expressed genes, we find it remarkable that so few deCOGs were recovered, and moreover that these gene sets are predominantly cytoskeletal and structural, rather than those genes classically involved in patterning and morphogenesis.

We therefore believe that our results add to a growing body of literature suggesting that the transcriptional components of regeneration are dissimilar across major animal clades. We note that the non-homology of animal regeneration at the transcriptional level does not negate the value of comparative studies across diverse taxa. Perhaps animal regeneration is homologous at another level of biological hierarchy (e.g. cell type regulation, tissue coordination), and the molecular logic coordinating this process evolved in an *ad hoc* manner across tissues and organisms. In this scenario, how conserved processes could be regulated by different molecular machinery would be the great challenge going forward. Alternatively, our results could signify true evolutionary convergence, in which case dozens—perhaps hundreds—of animal lineages have independently evolved solutions to bodily damage with varying degrees of success. Such a scenario puts a greater emphasis on natural selection driving regenerative capabilities, as opposed to such abilities being lost to genetic drift or countervailing selective forces. Given the apparent advantages of regeneration, how and why natural selection drives this trait in specific lineages is an interesting problem to study. Future studies across diverse animals will help to shed light on this question, and distinguish between the competing paradigms explaining the molecular and cellular mechanisms underlying regeneration.

## MATERIALS AND METHODS

### Data accessibility

The core code used to collect and analyze the RNA-Seq datasets is available through GitHub at https://github.com/nsierra1/RNAseq_pValueAggregation. Additional Files necessary for downstream analyses are avaialbe through Harvard Dataverse at https://doi.org/10.7910/DVN/LZK9DR.

### Transcriptome Collection

For the axolotl (*Ambystoma mexicanum*), a transcriptome was downloaded from the Broad Institute’s Axolotl Transcriptome Project (https://portals.broadinstitute.org/axolotlomics/; File: “Axolotl.Trinity.CellReports2017.fasta.gz”). For the planarian (*Schmidtea mediterranea*), a transcriptome was obtained from SmedGD (http://smedgd.stowers.org/; File: “SmedSxl Genome Annotations version 4.0 Predicted Nucleotide FASTA”). For the sea anemone (*Nematostella vectensis*) a transcriptome was downloaded from NCBI (BioProjects: PRJNA19965, PRJNA12581; File: “GCF_000209225.1_ASM20922v1_rna.fna”). For the sea cucumber (*Apostichopus japonicus*), reference isotigs were downloaded from the relevant paper (20) (NCBI accession: GSE44995; File: “GSE44995_Reference_assembled_isotig_seq.fna.gz”). For the sea sponge (*Halisarca caerulea*) the transcriptome was downloaded from the Figshare link provided in the original paper (File: “Halisarca_REF_trinity.fasta.zip”) (17). For the zebrafish (*Danio rerio*), all predicted cDNAs were downloaded from ENSEMBL release-89 (file: “GRCz10.cdna.all.fa”). The genes from these transcriptomes were converted into proteins using Transdecoder v5.0.2 (33), and are provided in **Additional File 2**.

### Read Collection and Mapping

RNA-Seq reads were downloaded from the NCBI Sequence Read Archive (SRA) using the “fastq-dump” program in the SRA Toolkit (https://www.ncbi.nlm.nih.gov/sra). **Table S3** provides a list of SRA IDs. The RNA-Seq reads were aligned to the relevant transcriptomes using HISAT-2 (34) and transcript abundances were quantified using RSEM v1.3.0 (35). The resulting RSEM quantifications are provided in **Table S3**, and the commands used to execute RSEM are reproduced in Additional File 1, part 0.1.

### Ortholog identification

The proteins determined by the transcripts from the six analyzed datasets were grouped into orthologous “gene sets” using the clustering algorithm OrthoFinder (23). The results of orthofinder analysis are provided in **Table S1**. All orthogroups are provided in **Additional File 1, part 1**. The resulting raw count matrices from RSEM were analyzed using EdgeR (36). We chose EdgeR because of its ability to accept a user-defined squareroot-dispersion value for studies that lack biological replication. The axolotl, cucumber, and sponge datasets lack biological replicates, making it impossible to estimate gene variance within samples. To deal with this shortcoming, we used EdgeR to see how various values for the biological coefficient of variation (BCV) impacted the number of differentially expressed genes. According to the EdgeR manual, typical values for BCV range from 0.4 for human data to 0.1 for genetically identical model organisms. We therefore tested a variety of BCV values within this space; the results are shown in **Figure S5**. Multidimensional scaling plots of BCV distances for samples with biological replicates are shown in **Figure S6**. We chose the lowest value for the squareroot-dispersion (0.1), in part because this allowed for the largest number of differentially expressed genes, and also because the spread of differentially expressed genes at various fold-change cutoffs behaved most similarly to datasets with biological replicates at this value (**Figure S5**). EdgeR was used to perform comparisons between adjacent time points. If a “wild-type” sample was included in the study, it was treated as equivalent to “time 0.” An example of the R code used to execute EdgeR is reproduced in the **Additional File 1, parts 0.2-0.3**. The resulting p-values and log count-per-million values were used for downstream aggregation of p-values and are also provided as Additional File 3.

**Figure S5.**
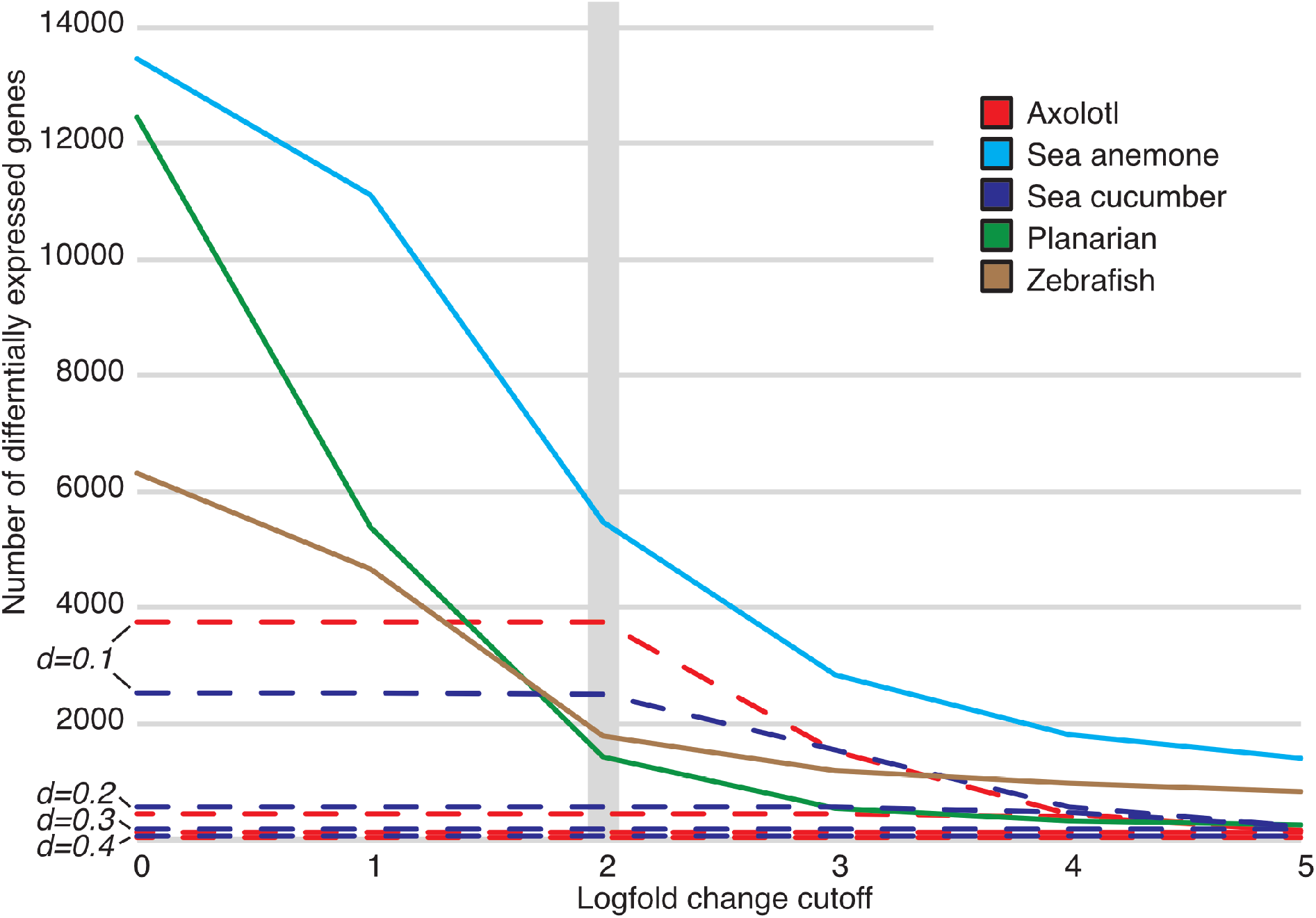
Impact of BCV values (denoted as “d”) on the number of differentially expressed genes in datasets lacking biological replication. The 2-fold change is noted with a grey bar; this is the standard logfold change cutoff for defining differentially expressed genes in RNA-Seq studies.

**Figure S6.**
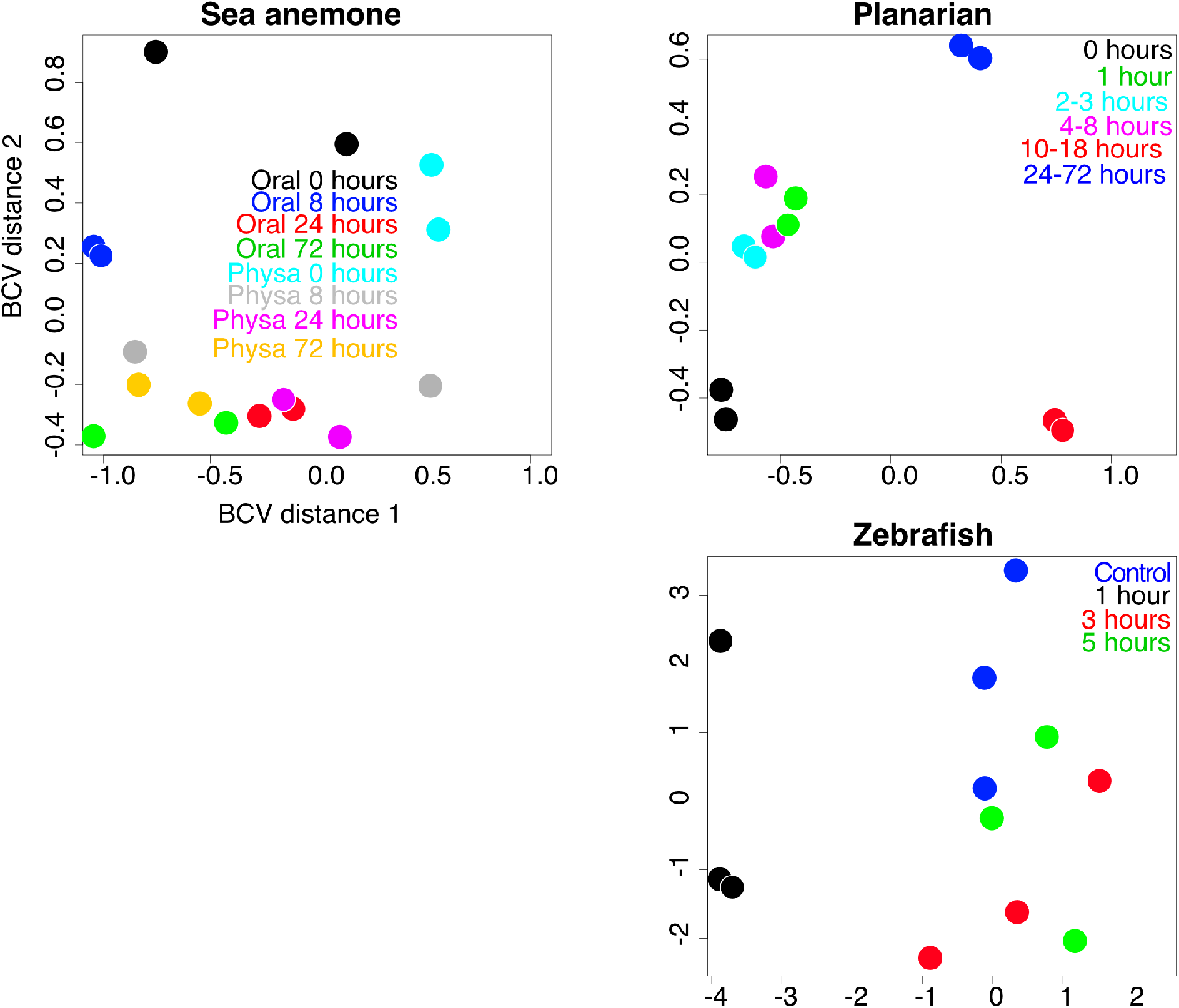
Multidimensional scaling plots of BCV distances between gene expression profiles for datasets containing biological replicates.

### p-value Aggregation

Aggregation of the p-values produced by EdgeR was based on methods described in Yi *et al.* (16). The method treated each p-value generated from adjacent time points for a given gene as an independent significance test of the null hypothesis that the broader COG was not differentially expressed. A resulting test of the *uniformity* for the set of p-values determines whether there is evidence that the COGs were not all unperturbed. Mathematically, the appropriate test statistic for uniformity can be computed from the sum of inverse cumulative distribution function with p-values and raw read counts as inputs. The result of this process is a table with entries corresponding to taxon-ortholog group pairs, and an associated aggregated p-value.

### False Discovery Rate Correction

Because each taxon has hundreds to thousands of distinct COGs, individual significance testing will result in many false positives. To ameliorate this, we perform the Benjamini-Hochberg procedure to adjust p-values for false discovery rate. The p-values were adjusted based on the total number of COGs such that no more than a constant fraction were likely to be false discoveries. These adjusted p-values were used for significance testing, and result in a list of ortholog groups corresponding to genes that likely to be differentially expressed during regeneration.

### Intersection Analysis

The final step was to derive a list of deCOGs shared across datasets. We originally attempted to do this by significance testing but found that numerical issues stemming from small p-values biased our tests such that a single p-value very close to 0 would yield a positive result, even if only one taxon showed strong results for that ortholog group. To avoid this problem, we used instead intersection analysis, looking at the presence/absence of deCOGs across datasets. This intersection method is less statistically rigorous but has the advantage of being robust to bias from small p-values.

### Correlation Plots and Venn Diagram

Overlap of COGs across taxa was visualized using correlation matrices and an Edwards Venn Diagram. A binary presence/absence table for each COG was modified from the output of OrthoFinder (provided in **Additional File 1, part 2.1**). A second table focused on presence/absence of deCOGs (**Additional File 1, part 2.2**). These tables were used to generate the correlation plots in **Figure 2** with the corrplot R library. Commands for generating the plots are provided in Additional File 1, part 2.3. The table of deCOGs was used to create an Edwards Venn Diagram using InteractiVenn^34^.

### Phylogenetic Assignment of Gene Families

Ideally, the evolutionary origin of each deCOG would be determined using a phylogenetically-informed clustering analysis such as OrthoFinder. Unfortunately taking such an approach at a eukaryote-wide scale is, for the time being, computationally prohibitive. Instead, we performed a series of BLAST queries and used sequence similarity of protein sequences to assign a phyletic origin for each COG.

Firstly, Uniprot Swissprot datasets were downloaded from www.Uniprot.com using the following queries:

1. Eukaryote (non-animal) dataset: *“NOT taxonomy:”Metazoa [33208]” AND reviewed:yes”*
2. Early animal dataset: *“taxonomy:”Metazoa [33208]”NOT taxonomy:”Bilateria [33213]” AND reviewed:yes”*
3. Bilaterian invertebrate dataset: *“taxonomy:”Bilateria [33213]” NOT taxonomy:”Vertebrata [7742]” AND reviewed:yes”*

Each of these datasets was turned into a BLAST database using the *makeblastdb* command. Our query COGs were the 2,770 deCOGs present in both the zebrafish and axolotl (see **Figure 3** of the main text), which also encompassed all deCOGs at broader evolutionary scales (i.e. the deCOGs shared by all vertebrates necessarily includes all deCOGs shared by deuterostomes, and so on). All protein sequences from these 2,770 deCOGs were collected and formatted into a query fasta file.

With the production of our query and database files, we proceeded with an iterative process of BLAST analyses. All proteins from the 2,770 deCOGs were queried against the “Eukaryote” database using BLASTp (command: *blastp -query Query_Proteins.fasta -db Eukaryote_Dataset - outfmt 6 -evalue 10e-5 -max_target_seqs 1 -num_threads 4 -out Results.txt*). If one or more queries had a hit, the entire deCOG was considered a “eukaryote novelty”. Proteins in the deCOGs that did not match anything in the “Eukaryote” database were used as the query sequences for the next BLASTp analysis against the “Early animal” database. Since sponges and other early-branching animals are poorly represented in Uniprot, any deCOG that had no match in the “Eukaryote” database and included at least one sponge protein was automatically designated as an “animal novelty,” regardless of whether or not it had a BLAST hit in the “Early animal” database. This process was repeated until all deCOGs were assigned a phyletic origin. A summary of these results is provided in **Additional File 1, part 6**.

### Enrichment analysis of deCOGs

Our comparison between all six taxa resulted in 160 deCOGs. We also examined the impact of individual taxa on the deCOG list by re-running the analysis with one organism excluded. Zebrafish (*Danio*) gene IDs from the resulting deCOGs were collected from each analysis, and are provided in **Additional File 1, part 3**. We restricted enrichment analysis to zebrafish genes that had at least one uncorrected (raw) p-value less than 0.01 from the original EdgeR analysis (**Additional File 1, part 0.2-0.3)**.

DAVID enrichment analysis was performed on the server (https://david.ncifcrf.gov). Zebrafish gene IDs were submitted using the “ENSEMBL_TRANSCRIPT_ID” identifier and a “Background” list type. STRING enrichment analysis requires a list of protein IDs, so the zebrafish transcripts were converted into protein identifiers using UniProt’s “Retrieve/ID mapping” function (https://www.uniprot.org/uploadlists/). The resulting IDs are provided in Additional File 1, part 3. These IDs were submitted to the STRING server for enrichment analysis (https://string-db.org). For both analyses, we restricted our study to conserved KEGG pathways. The full results of these analyses are provided in **Additional File 1, part 4**.

### Analysis of gene trees

In this paper, we examined the coverage of deCOGs in the KEGG stem cell pluripotency network (**Figure 5**). For genes present in all 6 datasets, we went back to the Orthofinder data to determine how gene families were organized into COGs, and which genes within those COGs were differentially expressed. Species-tree corrected gene trees were collected from the Orthofinder output. These trees were manually annotated to include gene names (based on zebrafish IDs) and whether or not genes were differentially expressed (smallest uncorrected p-value < 0.01 from EdgeR output). **Figure S4** shows the gene tree for *activin* and *bmp4* constructed using this method. The other trees were too large to illustrate as legible figures, but the tree in **Figure S4** and all additional, annotated trees are provided in newick format in Additional File 1, part 7.

## SUPPLEMENTAL TABLES

**Table S1.** Basic statistics of OrthoFinder analysis

|  |  |
| --- | --- |
| Number of genes | 538,991 |
| Number of genes in orthogroups | 266,324 |
| Number of unassigned genes | 272,667 |
| Percentage of genes in orthogroups | 49.4 |
| Number of orthogroups | 16,116 |
| Number of species-specific orthogroups | 1,447 |
| Number of genes in species-specific orthogroups | 18,356 |
| Percentage of genes in species-specific orthogroups | 3.4 |
| Mean orthogroup size | 16.5 |
| Median orthogroup size | 8 |
| Number of orthogroups with all species present | 2,287 |
| Number of single-copy orthogroups | 15 |

**Table S2:**
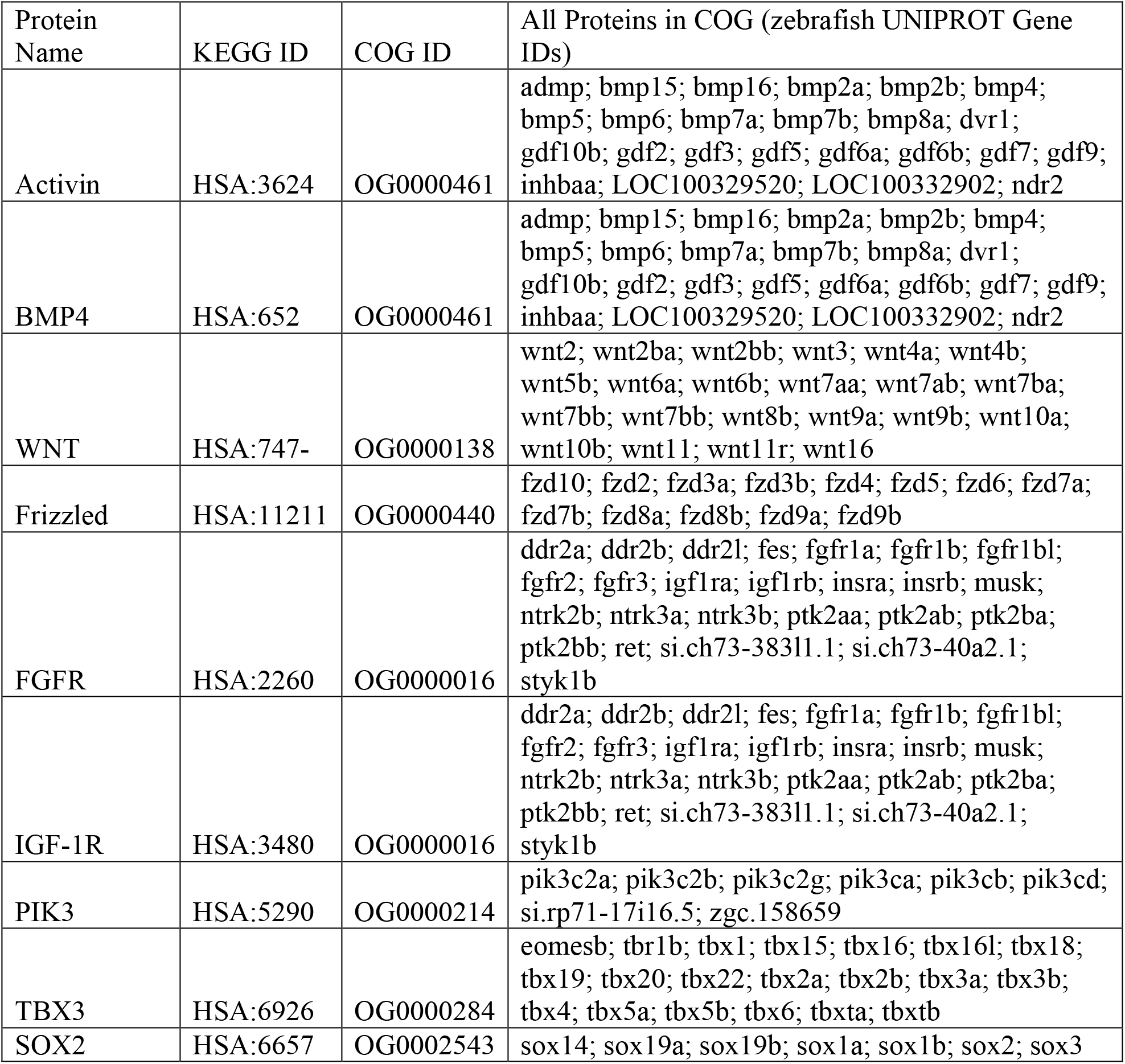
Paralogous zebrafish genes included in each conserved deCOG from Figure 5 of the main text.

**Table S3:** Alignment Statistics for RNA-Seq Data

| **Taxon** | **Dataset** | **Reads**<br>**Processed** | **Aligned**<br>**Reads** | **Unaligned**<br>**Reads** | **Suppressed**<br>**Reads** |
| --- | --- | --- | --- | --- | --- |
| Axolotl | 100309-lane1 | 16974066 | 77.02% | 18.69% | 4.29% |
|  | 100309-lane2 | 18631612 | 82.92% | 13.88% | 3.21% |
|  | 100309-lane3 | 18865464 | 87.02% | 11.31% | 1.68% |
|  | 100309-lane4 | 19355559 | 83.64% | 13.31% | 3.05% |
|  | 100309-lane5 | 19275194 | 85.61% | 12.68% | 1.71% |
|  | 100309-lane6 | 19450380 | 85.05% | 13.48% | 1.47% |
|  | 100309-lane7 | 18682600 | 78.67% | 17.89% | 3.44% |
| Planarian | ERR032066_1 | 24250265 | 43.21% | 56.79% | 0% |
|  | ERR032066_2 | 10333407 | 42.61% | 57.39% | 0% |
|  | ERR032067_1 | 24874216 | 44.63% | 55.37% | 0% |
|  | ERR032067_2 | 24874216 | 44.00% | 56.00% | 0% |
|  | ERR032068_1 | 20600012 | 44.04% | 55.96% | 0% |
|  | ERR032068_2 | 20600012 | 43.08% | 56.92% | 0% |
|  | ERR032069_1 | 24298493 | 29.36% | 70.64% | 0% |
|  | ERR032069_2 | 24298493 | 28.95% | 71.05% | 0% |
|  | ERR032070_1 | 28837238 | 28.66% | 71.33% | 0% |
|  | ERR032070_2 | 28837238 | 27.91% | 72.09% | 0% |
|  | ERR032071_1 | 23720712 | 45.89% | 54.11% | 0% |
|  | ERR032071_2 | 23720712 | 45.15% | 54.85% | 0% |
| Sea anemone | SRR3938202 | 34014046 | 65.25% | 34.75% | 0% |
|  | SRR3938203 | 28964964 | 63.49% | 36.51% | 0% |
|  | SRR3938286 | 21414087 | 63.05% | 36.95% | 0% |
|  | SRR3938287 | 13136751 | 63.29% | 36.71% | 0% |
|  | SRR3938288 | 40894772 | 59.95% | 40.05% | 0% |
|  | SRR3938289 | 32209584 | 62.39% | 37.61% | 0% |
|  | SRR3938290 | 11753545 | 59.86% | 40.14% | 0% |
|  | SRR3938291 | 19568370 | 64.77% | 35.23% | 0% |
|  | SRR3938293 | 15266653 | 64.28% | 35.72% | 0% |
|  | SRR3938294 | 17017293 | 61.84% | 38.16% | 0% |
|  | SRR3938297 | 26530337 | 64.06% | 35.94% | 0% |
|  | SRR3938298 | 23053972 | 63.36% | 36.64% | 0% |
|  | SRR3938299 | 21432871 | 60.17% | 39.83% | 0% |
|  | SRR3938300 | 43291231 | 59.66% | 40.34% | 0% |
|  | SRR3938303 | 16006522 | 61.86% | 38.13% | 0% |
|  | SRR3938304 | 11318128 | 58.99% | 41.00% | 0% |
| Sea cucumber | SRR771602 | 4871221 | 55.45% | 44.55% | 0.01% |
|  | SRR771606 | 5032070 | 52.56% | 47.43% | 0.01% |
|  | SRR771605 | 4729107 | 51.30% | 48.69% | 0.01% |
|  | SRR771604 | 4879963 | 56.53% | 43.47% | 0.01% |
|  | SRR771603 | 4716678 | 54.72% | 45.27% | 0.01% |
| Sea sponge | SRR5863988 | 9003557 | 74.68% | 25.12% | 0.20% |
|  | SRR5863987 | 9351918 | 49.98% | 49.63% | 0.39% |
|  | SRR5234759 | 15749742 | 52.35% | 47.34% | 0.31% |
| Zebrafish | SRR1205171 | 37130242 | 61.26% | 38.49% | 0% |
|  | SRR1205170 | 32103088 | 63.52% | 36.18% | 0.30% |
|  | SRR1205169 | 38057998 | 63.25% | 36.45% | 0.30% |
|  | SRR1205165 | 28133154 | 60.32% | 39.40% | 0.28% |
|  | SRR1205164 | 28183558 | 64.43% | 35.28% | 0.29% |
|  | SRR1205163 | 27285102 | 57.78% | 41.97% | 0.25% |
|  | SRR1205162 | 40651710 | 64.07% | 35.66% | 0.27% |
|  | SRR1205161 | 31246933 | 59.53% | 40.22% | 0.25% |
|  | SRR1205160 | 33781001 | 63.24% | 36.50% | 0.26% |
|  | SRR1205159 | 32238384 | 60.30% | 39.47% | 0.23% |
|  | SRR1205157 | 37460444 | 67.02% | 32.72% | 0.26% |
|  | SRR1205158 | 33073599 | 53.06% | 46.74% | 0.20% |
|  | SRR1205157 | 37460444 | 67.02% | 32.72% | 0.26% |

## Notes

https://github.com/nsierra1/RNAseq_pValueAggregation

https://doi.org/10.7910/DVN/LZK9DR

### Competing Interest Statement

The authors have declared no competing interest.

